# R-based method for quantitative analysis of biofilm thickness by using Confocal Laser Scanning Microscopy

**DOI:** 10.1101/2022.02.13.480190

**Authors:** Hanna Marianne Frühauf, Markus Stöckl, Dirk Holtmann

## Abstract

Microscopy is mostly the method of choice to analyse biofilms. Due to the high local heterogeneity of biofilms, single and punctual analyses only give an incomplete insight into the local distribution of biofilms. In order to retrieve statistically significant results a quantitative method for biofilm thickness measurements was developed based on confocal laser scanning microscopy and the programming language R. The R-script allows the analysis of large image volumes with little hands-on work and outputs statistical information on homogeneity of surface coverage and overall biofilm thickness. The applicability of the script was shown in microbial fuel cell experiments. It was found that *G. sulfurreducens* responds differently to poised anodes of different material so that the optimum potential for MFC on poised ITO anodes had to be identified with respect to maximum current density, biofilm thickness and MFC start-up time. Thereby, a positive correlation between current density and biofilm thickness was found, but with no direct link to the applied potential. The optimum potential turned out to be +0.1 V vs SHE. The script proved to be a valuable stand-alone tool to quantify biofilm thickness in a statistically valid manner, which is required in many studies.

**Practical application:** Biofilm communities are ubiquitous. They can be found in every habitat in which water, nutrients and a colonisable surface are present. Depending on the surface, biofilms can cause economic losses due to bio-corrosion (pipelines and ship walls are prominent examples) or are a severe threat to human health when important medical devices or body tissues are colonised ^[1]^. Desirable biofilms are catalytic biofilms how they are used in bioelectrochemical production processes, for example. In all cases, quantitative and qualitative biofilm analysis is necessary in order to prevent or promote biofilm formation. In bioelectrochemistry quantitative biofilm analysis is essential to link productivity (current or chemicals) with biomass deposited on the electrode. Microscopic analysis (e.g. with CLSM) of stained biofilms allows the recording of high volumes of image data but often image analysis then remains at a qualitative stage. In terms of biofilm thickness determination this limits analysis to an estimated thickness of a small amount of images, mostly. The presented R-script allows the calculation of biofilm thickness based on a larger amount of image sets and allows conclusions on the homogeneity of biofilm coverage on the electrode surface. The script is a stand-alone tool if only biofilm thickness should be determined and does not require any image segmentation or processing.

## 1 Introduction

Biofilms appear ubiquitously in non-natural habitats like heat exchangers, pipes, hoses, medical implants and catheters but are also applied in catalytic processes. Heterogeneity in biofilm growth plays an important role in controlling biofouling, biocorrosion or application of antibiotics as well as improving production processes with catalytic biofilms. From an engineering point of view, one of the first steps required to assess the statistical significance (or possibly insignificance) of heterogeneity in biofilm growth and its variability ^[2]^.

Confocal laser scanning microscopy (CLSM) is a non-destructive and non-invasive method with the capability to provide three-dimensional images of biofilms. The analysis of the obtained CLSM data is very often rather qualitative than quantitative and based on a subjective visual image inspection. This approach is however not feasible when large quantities of data have to be analysed, which is often necessary to ensure the significance of the outcome of the CLSM measurements ^[3]^. In order to perform a quantitative analysis an R-based method was developed and evaluated in a bioelectrochemical process with the biofilm-forming and electroactive organism *G. sulfurreducens*.

*G. sulfurreducens* is a model organism in microbial fuel cell (MFC) research and known for its good ability to form biofilms on the electrode and high current densities produced from complete acetate oxidation ^[4]^. Mostly, carbon-based materials are used as anode material because of their good biocompatibility, corrosion-resistance and cost-effectiveness ^[5]^. Especially for spectroscopic methods in connection with biofilm analysis transparent anodes are required like indium-tin oxide (ITO) or gold sputtered glass slides (e.g. in ^[6]^, ^[7]^, ^[8]^). Most lab-scale MFC are operated with poised anodes to drive *G. sulfurreducens* extracellular electron transfer. Thereby, the more positive the potential, the higher the energy supplied to the system. Direct electron transfer between *G. sulfurreducens* and the anode is based on a relay of outer membrane cytochromes (Omc) and conductive pili ^[9]^ ^[10]^ ^[11]^ ^[12]^. This network allows the formation of conductive biofilms in which also the outer cell layers are electrically “wired” to the anode ^[13]^ ^[14]^. It was shown that the applied potential influences the type of Omc active in electron transport as well as the composition of the extracellular polymeric substances present in the biofilm ^[15]^ ^[16]^ ^[17]^. What has been little discussed so far, is the interpolated effect of electrode material and applied potential. *G. sulfurreducens* MFC were conducted at different potentials on ITO electrodes to determine a potential which optimises the MFC performance indicators maximum current density, start-up phase and biofilm thickness. The latter characteristic is often only evaluated qualitatively when CLSM is used for analysis as manual thickness determination from CLSM images is laborious and time-consuming. CLSM is the method of choice as it allows rapid qualitative evaluation of biofilm heterogeneity, electrode coverage and cell viability (if appropriate staining methods are used). With fluorescent strains biofilms can also be analysed in vivo, i.e. during the experiment, which gives insights into MFC start-up time for example. Other methods for thickness determination require the knowledge of the biofilm refractive index (analysis with light microscopy) ^[18]^, additional preparatory steps like cryo-cut ^[2]^ or special equipment, unusual in bioelectrochemical applications (low-load compression testing in ^[19]^).

The optimisation of *G. sulfurreducens* MFC performance on ITO electrodes was used as a proof-of-principle for a quantitative method that measures biofilm thickness based on CLSM imaging of fluorescently stained biofilms and statistical evaluation with the programming language R ^[20]^. It is a stand-alone method that does not require image segmentation or filtering and allows statistical analysis of a high volume of biofilm images with little hands-on work. Explanation and evaluation of the R-script is central to this publication, and the optimisation of MFC performance on ITO electrodes should be seen as an exemplary application rather than a profound explanation of the biological phenomenon.

## 2 Materials and Methods

### 2.1 Cultivation of *G. sulfurreducens*

*G. sulfurreducens* strain PCA (DSM 12127) was obtained from DSMZ (German collection of Microorganisms and Cell Cultures GmbH, Braunschweig, Germany). All cultivations were done anaerobically in serum flasks sealed with a butyl septum. Flasks were incubated shaking at 30 °C and 180 rpm. Growth medium was DSM826 as recommended by DSMZ, but with 30 mM Na_2_-fumarate instead of 50 mM as soluble electron acceptor. Cultivation conditions were also described in ^[21]^.

### 2.2 Microbial fuel cell experiments with *G. sulfurreducens*

MFC experiments were carried out in a modified electrochemical H-cell, as described in ^[21]^. In this setup the WE is flanged to the WE chamber from one side which ensures a stable fixation in the reactor and thereby stable flow conditions for bacterial adhesion and biofilm growth. ITO coated glass was used as WE (30mm × 30mm × 1.1mm; resistance 20 Ω cm^−2^; Präzisions Glas & Optik GmbH, Iserlohn, Germany) with graphite paper for contacting (0.13 mm thin, type RCT®-DKA-SBGR; RCT Reichelt Chemietechnik GmbH + Co., Heidelberg, Germany) and graphite as counter electrode (Bipolar plate, Eisenhuth GmbH & Co. KG, Osterode am Harz, Germany). Experiments were conducted at constant potential by using IPS (Elektronik GmbH & Co KG, Münster, Germany) and Gamry potentiostats (Reference600 and Interface1000; Gamry Instruments, Warminster, PA) with Ag/AgCl/KCl_sat_ reference electrode (= SHE - 0.2 V; Sensortechnik Meinsberg, Waldheim, Germany) that was placed in a Haber-Luggin-capillary filled with KCl_sat_. Both chambers were filled with growth medium lacking the soluble electron acceptor fumarate.

### 2.3 CLSM analysis

Immediately after an experiment was aborted, biofilms were stained with the fluorescent dyes SYTO^TM^9 Green Fluorescent Nucleic Acid Stain (Syto9) and Propidium iodide (PI; both InvitrogenTM,Waltham MA, USA). Stock solutions in 0.125 M phosphate buffer were 6 mM Syto9 and 30 mM PI. Dyes were mixed 1:1 in a final volume of 400 μl per biofilm (stained area 5 cm^2^). Electrodes with biofilms were carefully detached from the WE chamber, placed in a petri dish (5 cm diameter), overlaid with 400 μl staining solution and incubated for 15 min in the dark. Afterwards, staining solution was removed and the biofilm rinsed three times with 0.125 M phosphate buffer, the fourth buffer volume remained on the biofilm to avoid desiccation.

Microscopic images were acquired with a TCS SP8 CLSM (Leica Microsystems GmbH, Wetzlar, Germany) with galvanometric stage. In order to image the biofilm in a hydrated state and thereby retrieving a realistic value for biofilm thickness, a dip-in objective was used (HC APO UVIS CS2, 63x magnification, numerical aperture 0.9, refraction index 1.33) that allowed imaging while the biofilms were stored in a petri dish, filled with buffer. Laser intensity (OPSL 488) was set to 5% and an excitation beam splitter DD488/552 used with the PMT detector set to emission wavelengths between 500 and 545 nm (for the green channel) and the HyD between 615 and 788 nm (for the red channel). The basis to determine biofilm thickness were z-images of a 185 μm wide area taken on ten randomly chosen sections on the biofilm. For optimum analysis laser intensity should be chosen in a way that the biofilm is clearly distinguished but not overexposed.

### 2.4 Quantification of biofilm thickness using R

Ten regions of interest (ROI) were defined on the z-images, each with one tenth of the image width, i.e. 18.5 μm (shown in Figure S1). Fluorescence intensity was recorded along the length of each ROI and plotted against the z-axis, resulting in a histogram (Figure 1). The thickness of the biofilm was extracted from the range of the histogram in which fluorescence was above a certain threshold level that defined the background fluorescence (indicated by the dashed line in Figure 1). Histogram data were exported from the LAS X software (Version 3.5.5) as .csv files which were then input for an R-script with which biofilm thickness was calculated.

**Figure 1:**
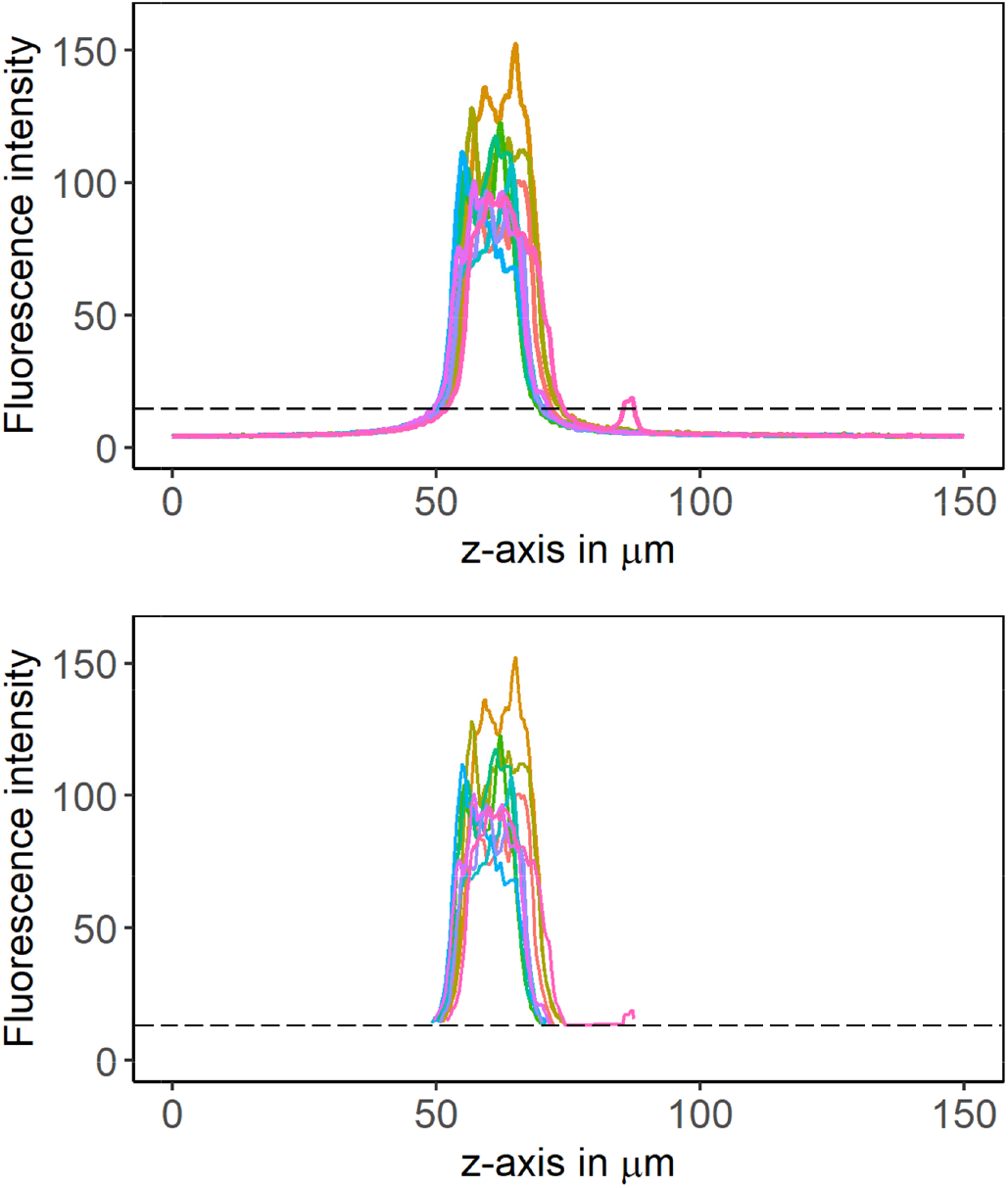
Fluorescence intensity of 10 ROI in one z-image are plotted against the z-dimension, before (top) and after (bottom) background subtraction. The dashed line indicates the threshold cut off. The area to the right of the maximum is biologically above the biofilm, i.e. buffer and planctonic cells. To the left is the electrode accordingly. In the shown example a fluorescence signal caused by planktonic cells could not be subtracted by the defined threshold, hence additional data treatment was needed so that biofilm thickness is not overestimated.

The R-script is shown in the SI and provided as .txt file along with this paper. The basic function of the script is explained in the following, with further details in the SI. The R-packages needed are data.table, reshape2, dplyr, (ggplot2 and cowplot for plotting). The .csv-files exported from LAS X must be converted to UTF-8 first, otherwise the R function *fread* is unable to read the files. This can be done for example with the online tool subtitletools, available on https://subtitletools.com/convert-text-files-to-utf8-online. The fluorescence intensity data (line profils) that were used in this study are available in “figshare” at https://doi.org/10.6084/m9.figshare.19165097.v1.

Pixels in the z-image that belong to the biofilm (and not to the background) are defined as those with intensity higher than 3-times the mean intensity of the background signal. All background pixels are replaced with NA, which is an “empty” value for the script. Since the background above the biofilm (buffer and planctonic cells) and below (electrode) have different background intensities the calculation is performed separately for left (first line; id < id_max) and right (first line; id > id_max) of the histogram's maximum (script line 36 – 40). That is why the information about the histogram peak was stored in the data column “id_max” before (line 35). Here the threshold is defined as 3-times mean background intensity but for higher background signal, i.e. background fluorescence of a different electrode material the threshold might need to be adapted (e.g. 2-times the background signal).

While the border between the electrode and the biofilm is defined as the point at which the intensity exceeds 3*mean(backgroundsignal), it is more difficult for the “End” of the biofilm (the transition between the biofilm and the buffer). Even though the biofilm is washed prior to CLSM analysis there are always cells floating above the biofilm that disturb the measurement. These fluorescent spots on the image can have signal intensity > 3-times background intensity and thereby falsify the definition of the biofilm end (see Figure 1). A rescue for this is to define the first occurring case in the histogram at which the intensity is < 3*mean(background) as the biofilm end. Since row numbers were added to the data table in the beginning (line 34) and all background pixels were eliminated from the table afterwards, the first case is the first gap in the consecutive row number column (with id > id_max, i.e. after maximum of the histogram). The gap can be detected by calculating the difference between entries of the row number column (line 68 - 73). Hence, the script “searches” for any difference other than 1 (line 74 - 76). While this solves the problem for the “End” of the biofilm, it might now overlook the “Beginning” of the biofilm if there is no gap in the row number. This is bypassed by doubling the first row of each histogram (line 64 - 67). The difference in rownumber is then 0 for these cases and thereby grabbed by the script. When the difference is calculated, the resulting data column is one entry short compared to the rownumber column, so an additional entry had to be added to the column (0 in this case) (line 70 -71).

## 3 Results and Discussion

*G. sulfurreducens* MFC were performed on ITO electrodes, poised at different potentials (−0.1, 0, +0.05, +0.1, +0.2, +0.3 V vs SHE) to identify the potential with the best response with respect to the performance parameters maximum current density, start-up phase and biofilm thickness. Results are shown in Table 1 (mean and SD of n = 3). Parameters related to current production were extracted from the potentiostat record and biofilm thickness determined with the developed R-script.

**Table 1.** Maximum current density, start-up time, biofilm thickness and CE for different potentials applied in MFC on ITO electrodes. Indicated are mean values ± SD (n = 3).

| Applied potential in V vs SHE | Maximum current density in $\mu\text{A cm}^{-2}$ | Start-up time in h | Biofilm thickness in $\mu\text{m}$ |
| --- | --- | --- | --- |
| -0.1 | $244 \pm 19\%$ | $20.4 \pm 20\%$ | $23 \pm 7$ |
| 0 | $352 \pm 24\%$ | $19.8 \pm 13\%$ | $35 \pm 6$ |
| +0.05 | $377 \pm 35\%$ | $20.0 \pm 26\%$ | $34 \pm 13$ |
| +0.1 | $399 \pm 24\%$ | $19.2 \pm 7\%$ | $38 \pm 7$ |
| +0.2 | $384 \pm 38\%$ | $22.3 \pm 4\%$ | $35 \pm 7$ |
| +0.3 | $417 \pm 20\%$ | $41.4 \pm 12\%$ | $29 \pm 7$ |

In order to assess any interdependence between the applied potential and the MFC performance indicators, Pearson correlation coefficients (r) were calculated, using the function *rcorr()* from the R-package *Hmisc*. Mathematically there was no linear correlation between the applied potential and any of the performance indicators but biofilm thickness correlated positively the maximum current density (r = 0.79, p < 0.0001), hence the thicker the biofilm, the higher the maximum current. The correlation is illustrated in Figure 2 which shows the maximum current density of each experiment plotted against the measured biofilm thickness. Noticeable is the high deviation for the triplicates (three experiments with identical potential applied but conducted on different days) in maximum current density, which is the highest for +0.05 and +0.2 V vs SHE applied (35, respectively 38%). Despite the deviation for the triplicates the positive correlation is still valid if all experiments are treated individually. As biofilm thickness is measured methodically at different areas on the electrode, each thickness measurement for whole electrodes is automatically linked to an estimate of the statistical uncertainty (shown as horizontal error bar in Figure 2). The width of the error bar indicates the homogeneity of thickness on the whole electrode area. The experiment polarised at +0.05 V vs SHE with the highest maximum current density for example shows a higher SD compared to the other electrodes (± 9 μM).

**Figure 2.**
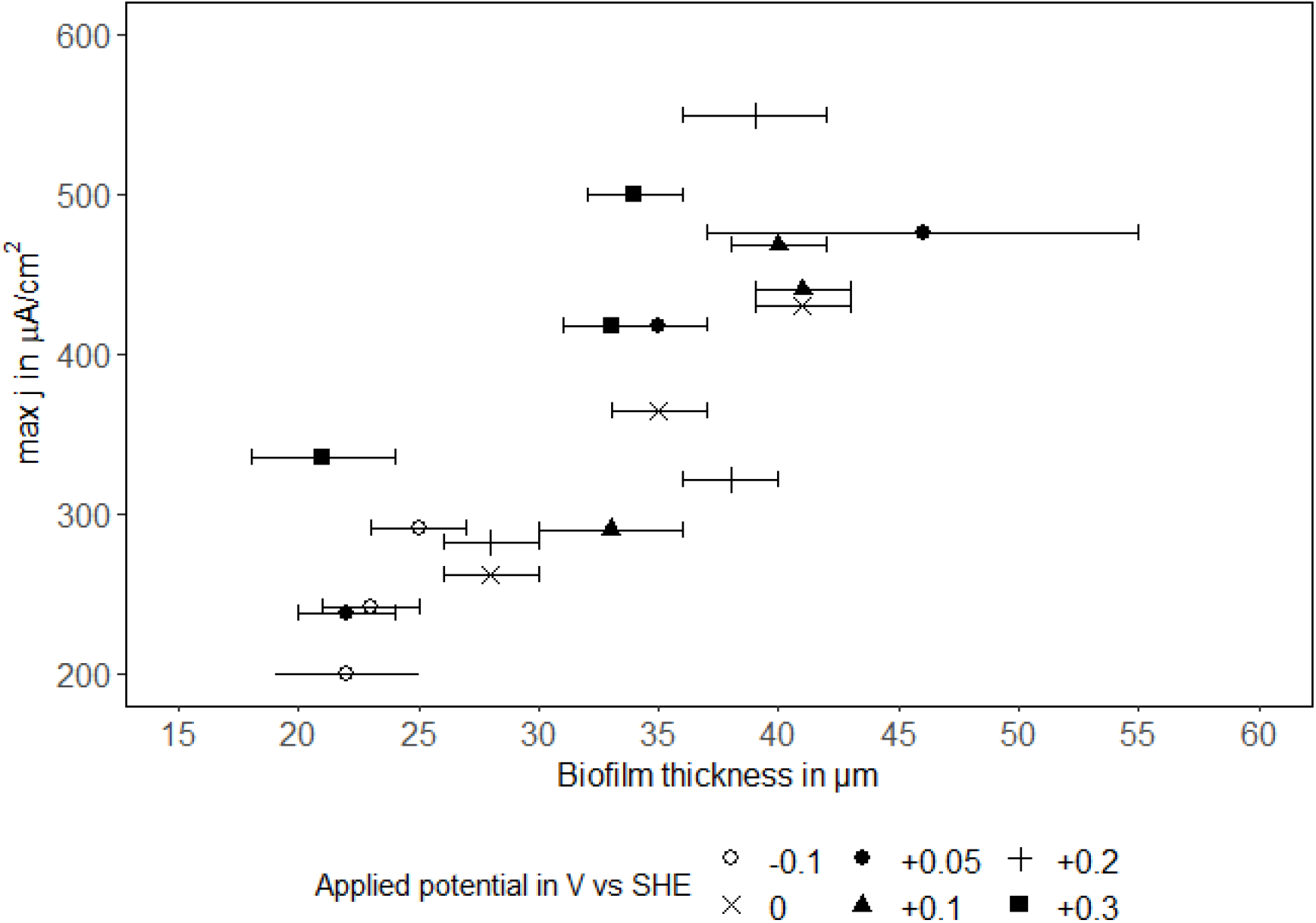
The maximum current density j is plotted against the mean biofilm thickness (n = 10) to illustrate the positive correlation. Horizontal error bars show SD of ten z-images analysed on each biofilm.

The reason for this can be seen in Figure 3 which displays the determined thickness for the electrode polarised to +0.05 V vs SHE with the highest deviation. Shown are boxplots of ten ROI assigned per image, with the corresponding position on the electrode on the x-axis. The thickness for 7, 8, 9 and 10 is significantly larger compared to the other images. The location of the images on the electrode are marked in the image insert and displays that in this experiment the lower part of the electrode was significantly more attractive for biofilm formation compared to the centre or the upper part. To be sure that this was not a methodical error, the results from the script were reviewed with the z-images and a qualitative assessment of the thickness by laying out a scale was done. The results from the script and the manual analysis differed by a mean of 7% (n = 10, SD = 2%) which verifies the results from the script as a true biological effect.

**Figure 3.**
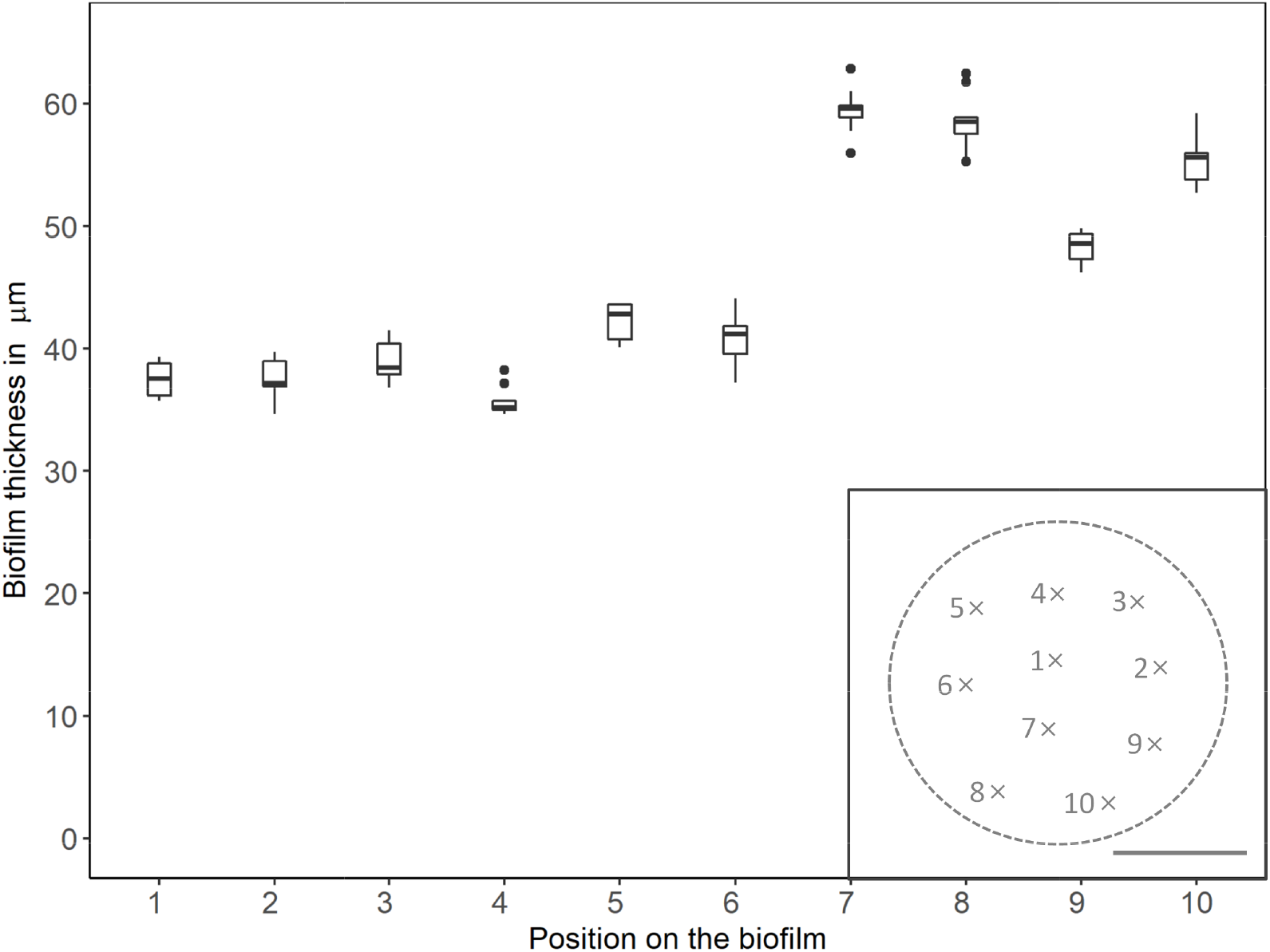
Calculated biofilm thickness is plotted with the corresponding location on the biofilm (position on x-axis, approximate position mapped on biofilm in insert). The scale bar in the insert is equivalent to 1 cm.

## 4 Concluding remarks

The described method allowed a statistically significant assessment of homogeneity of biofilm coverage on the electrode as well as the identification of a correlation between biofilm thickness and current density that did not depend directly on the applied potential. The most favourable potential on ITO anodes was +0.1 V vs SHE with a maximum current density of almost 400 μA cm^−2^ and 38 ± 7 μm biofilm thickness. Of course biofilm thickness determination can also be done manually but is less accurate and more time consuming, as well as the variation within one z-image cannot be included. For the script developed in this work no image correction has to be performed which simplifies the analysis additionally. The recently published software package “BiofilmQ” ^[22]^ also covers biofilm thickness calculation, among a vast number of other functions but might be oversized for some applications. Further “BiofilmQ” is based on Matlab, so if there is an existing R routine the stand-alone script from this work could easily be integrated. The presented open-access script can be applied to all experimental questions for which analysis of biofilm thickness and finally biofilm topology is scientifically or technically relevant.

## Supporting information

R-script

## Abbreviations

CLSM: confocal laser scanning microscopy
ITO: indium-tin oxide
MFC: microbial fuel cells

## Acknowledgements

The authors gratefully acknowledge the financial support from the German Federal Ministry of Education and Research (nos. 031A226). The authors have declared no conflicts of interest.

## Data availability

The data that support the findings of this study are openly available in “figshare” at https://doi.org/10.6084/m9.figshare.19165097.v1.

## References

[1] a) L. Hall-Stoodley, J. W. Costerton, P. Stoodley, Nature reviews microbiology 2004, 2, 95–108; b) W. Dou, D. Xu, T. Gu, Microbial Biotechnology 2021, 14, 803–805.

[2] R. Murga, P. S. Stewart, D. Daly, Biotechnology and bioengineering 1995, 45, 503–510.

[3] A. Heydorn, B. K. Ersbøll, M. Hentzer, M. R. Parsek, M. Givskov, S. Molin, Microbiology 2000, 146, 2409–2415.

[4] D. R. Bond, D. R. Lovley, Applied and environmental microbiology 2003, 69, 1548–1555.

[5] A. Baudler, I. Schmidt, M. Langner, A. Greiner, U. Schröder, Energy & Environmental Science 2015, 8, 2048–2055.

[6] L. Robuschi, J. P. Tomba, J. P. Busalmen, Journal of Electroanalytical Chemistry 2017, 793, 99–103.

[7] J. Song, D. Sasaki, K. Sasaki, S. Kato, A. Kondo, K. Hashimoto, S. Nakanishi, Process Biochemistry 2016, 51, 34–38.

[8] F. Scarabotti, L. Rago, K. Bühler, F. Harnisch, Bioelectrochemistry 2021, 140, 107752.

[9] K. P. Nevin, H. Richter, S. Covalla, J. Johnson, T. Woodard, A. Orloff, H. Jia, M. Zhang, D. Lovley, Environmental microbiology 2008, 10, 2505–2514.

[10] K. P. Nevin, B.-C. Kim, R. H. Glaven, J. P. Johnson, T. L. Woodard, B. A. Methé, R. J. DiDonato Jr, S. F. Covalla, A. E. Franks, A. Liu, PloS one 2009, 4, e5628.

[11] T. Mehta, M. V. Coppi, S. E. Childers, D. R. Lovley, Applied and environmental microbiology 2005, 71, 8634–8641.

[12] D. R. Bond, S. M. Strycharz-Glaven, L. M. Tender, C. I. Torres, ChemSusChem 2012, 5, 1099.

[13] a) G. Reguera, K. P. Nevin, J. S. Nicoll, S. F. Covalla, T. L. Woodard, D. R. Lovley, Applied and environmental microbiology 2006, 72, 7345–7348; b) G. Reguera, K. D. McCarthy, T. Mehta, J. S. Nicoll, M. T. Tuominen, D. R. Lovley, Nature 2005, 435, 1098–1101.

[14] X. Liu, D. J. Walker, S. S. Nonnenmann, D. Sun, D. R. Lovley, Mbio 2021, 12, e02209–02221.

[15] D. B. Li, J. Li, D. F. Liu, X. Ma, L. Cheng, W. W. Li, C. Qian, Y. Mu, H. Q. Yu, Biotechnology and bioengineering 2019, 116, 961–971.

[16] D.-F. Liu, W.-W. Li, Current Opinion in Chemical Biology 2020, 59, 140–146.

[17] G. Yang, L. Huang, Z. Yu, X. Liu, S. Chen, J. Zeng, S. Zhou, L. Zhuang, Water research 2019, 159, 294–301.

[18] R. Bakke, P. Olsson, Journal of Microbiological Methods 1986, 5, 93–98.

[19] E. Paramonova, E. D. De Jong, B. P. Krom, H. C. Van der Mei, H. J. Busscher, P. K. Sharma, Applied and environmental microbiology 2007, 73, 7023–7028.

[20] R. C. Team, 2013.

[21] M. Stöckl, N. C. Teubner, D. Holtmann, K.-M. Mangold, W. Sand, ACS applied materials & interfaces 2019, 11, 8961–8968.

[22] R. Hartmann, H. Jeckel, E. Jelli, P. K. Singh, S. Vaidya, M. Bayer, D. K. Rode, L. Vidakovic, F. Díaz-Pascual, J. C. Fong, Nature microbiology 2021, 6, 151–156.

