## Supplementary information for "R-based method for quantitative analysis of biofilm thickness by using Confocal Laser Scanning Microscopy"

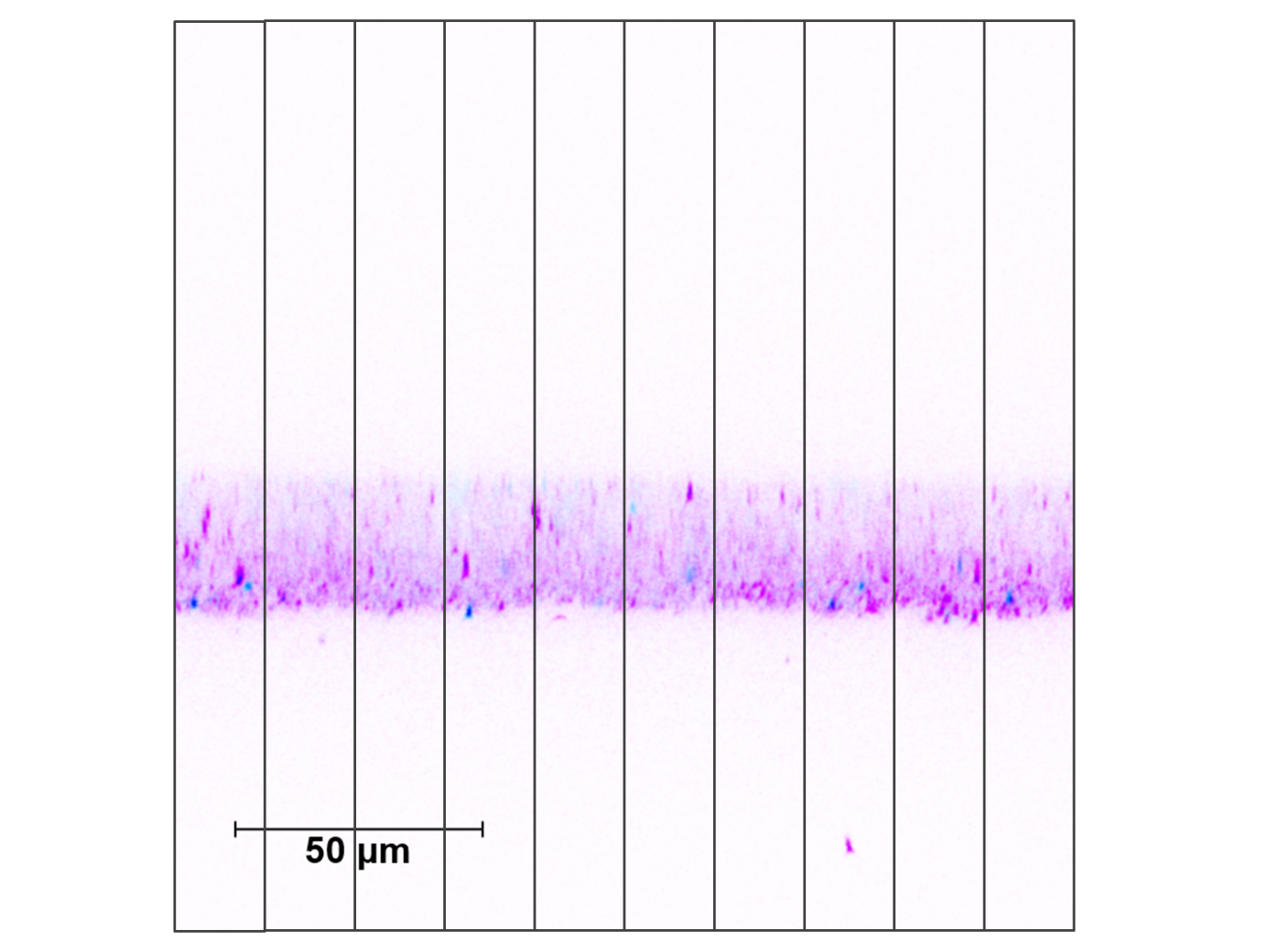

**Figure S1.** Ten regions of interest were defined on the z-images, each with one tenth of the image width, i.e. 18.5 μm. Fluorescence intensity is recorded along the length of each ROI and plotted against the z-dimension, resulting in a histogram (Figure 1).

**Table S1**. Commented R-script; plain (non-formatted) version of the script is also available for download.

| 1  2  3  4  5  6  7  8  9  10  11  12  13  14  15  16  17  18  19  20  21  22  23  24  25  26  27  28  29  30  31  32  33  34  35  36  37  38  39  40  41  42  43  44  45  46  47  48  49  50  51  52  53  54  55  56  57  58  59  60  61  62  63  64  65  66  67  68  69  70  71  72  73  74  75  76  77  78  79  80  81  82  83  84  85  86  87  88  89  90  91  92  93  94  95  96  97  98  99  100  101  102  103  104  105  106  107  108  109  110  111  112  113  114  115  116  117  118  119  120  121  122  123  124  125  126  127  128 | # Load packages  library("ggplot2")  library("reshape2")  library("dplyr")  library("cowplot")  library("data.table")  # First, a list of all filenames is extracted from the folder in which the files are stored. The file ending .csv is used as the identifier (pattern)  filenames 🡨 list.files("CLSM/Line profils", pattern="*.csv", full.names=TRUE)  # All files are read with the function fread and stored in a list, combined with the file names  data 🡨 lapply(filenames, fread)  # There is a separate column in which z-axis coordinates are stored for each ROI separately but since it is identical for each ROI, the redundant columns can be deleted.  # The argument of seq() depends on the number of ROI defined in one image. seq(3,19,2) applies for 10 ROI in one image.  data 🡨 lapply(data, function(x) x[,-seq(3,19,2)])  # In "data" intensity values are arranged column-wise, which is then melted to a tidy data set with intensity values stored row-wise in "value" and the ROI names as identifier stored in "variable"  data_melt 🡨 lapply(data, function (x) melt(x, id = c("Axis [m]")))  # An additional identifier column with the last part of the file name is added to link data with the information which electrode was analysed. Therefore path before filenames is removed (keep expression after square break; attention – sub() leaves blank space before the name)  filenames_short 🡨 sub(".*]", "", filenames)  data_melt 🡨 mapply(cbind, data_melt, "file"=filenames_short, SIMPLIFY=F)  # For easier data handling the tables are converted to data frames  df_data_melt 🡨 do.call(rbind.data.frame, data_melt)  # Missing values are deleted from the data frames and the z-axis coordinates are converted from meter to micrometer  df_data_melt 🡨 df_data_melt[complete.cases(df_data_melt), ]  df_data_melt$file 🡨 as.factor(df_data_melt$file)  df_data_melt$`Axis [m]` 🡨 df_data_melt$`Axis [m]`*10^6  # For each file-ROI combination an extra column is added which indicates the row with the maximum fluorescence intensity, i.e. the peak of the histogram (id_max). In addition a row number is added (id). This is needed later to identify the upper and the lower boarder of the biofilm  df_data_melt 🡨 df_data_melt %>%  group_by(file, variable) %>%  mutate(id = row_number(),  id_max = id[value == max(value)])  # Replace background with NA (separately for left and right of the peak)  replace_background 🡨 df_data_melt %>%  group_by(file, variable) %>%  mutate(value = replace(value, id < id_max & value < 3*mean(head(value, n = 100)), NA),  value = replace(value, id > id_max & value < 3*mean(tail(value, n = 100)), NA))  # Delete NA from data frame  no_background 🡨 na.omit(replace_background)  # Plots of the data frames before and after subtracting the background signal to randomly check on the correctness of the threshold definition  minus_background 🡨 ggplot(subset(no_background, file == " 2_5 z.csv"), aes(`Axis [m]`, value)) +  geom_point(aes(colour = variable), size = 0.75) +  scale_x_continuous(limits = c(0, 200)) +  scale_y_continuous(limits = c(0, 200)) +  xlab(expression(paste("z-axis in ", mu, "m"))) +  ylab("Fluorescence intensity") +  plot.options.legend +  theme(legend.position = "none")  all 🡨 ggplot(subset(df_data_melt, file == " 5_2 z.csv"), aes(`Axis [m]`, value)) +  geom_point(aes(colour = variable), size = 0.75) +  scale_y_continuous(limits = c(0, 200)) +  scale_x_continuous(limits = c(0, 200)) +  xlab(expression(paste("z-axis in ", mu, "m"))) +  ylab("Fluorescence intensity") +  plot.options.legend +  theme(legend.position = "none")  plot_grid(all, minus_background, align = "v", nrow = 2)  #### Detailed information on second part of script can be found in the Material & Methods section  # Duplicates the first row of every file-variable combination  no_background2 🡨 no_background %>%  group_by(file, variable) %>%  slice(rep(1:n(), c(2, rep(1, each = n()-1))))  # calculates the difference in rownumber between consecutive rows  diff 🡨 diff(no_background2$id)  # adds an additional item to the end of the vector, number 0 in this case  diff[length(diff) + 1] 🡨 0  # adds the vector to the data frame  no_background2$diff 🡨 diff  # This step extracts the gaps resulting from the deleted background, by filtering all rows for which the vector "diff" is not "1" (i.e. non-consecutive rows)  all_gaps 🡨 subset(no_background2, diff != 1)  # For some images the background line left if the histogram (the "electrode-part") oscillates around values greater and smaller than 3*mean(background) which also causes gaps in the "diff"-vector but relevant to define the “End” of the biofilm are only values to the right of the maximum, i.e. id > id_max. The "End" of the biofilm is then the first value for each image-ROI (= file-variable) combination stored in the “Axis” column of the all_gaps2 data frame.  all_gaps2 🡨 all_gaps %>%  group_by(file, variable) %>%  subset(id > id_max)  end 🡨 all_gaps2 %>%  group_by(file, variable) %>%  summarise(end = `Axis [m]`[1])  # For the "Begin" of the biofilm in turn the last gap to the left of the maximum is relevant, i.e. id < id_max. The "Begin" of the biofilm is then the last value for each image-ROI (= file-variable) combination stored in the “Axis” column of the all_gaps_begin data frame.  all_gaps_begin 🡨 all_gaps %>%  group_by(file, variable) %>%  subset(id < id_max)  summary 🡨 all_gaps_begin %>%  group_by(file, variable) %>%  summarise(begin = tail(`Axis [m]`, 1))  # Values for the "End" are added to the data table in which the values for the "Begin" are already stored. Then, the biofilm thickness can be calculated as the difference between the z-axis coordinates for "End" and "Begin".  summary$end 🡨 end$end  summary$thickness 🡨 summary$end - summary$begin  # To summarise the results for each electrode, an electrode indicator is added to the table (there are 100 file-ROI combinations, therefore each electrode name is replicated 100 times).  summary$electrode 🡨 as.factor(rep(seq(1,6), each = 100))  # The result can be plotted split up to each image and ROI or summarised in a boxplot  ggplot(summary) +  geom_point(aes(x = file, y = thickness, colour = variable)) +  xlab("Filename") +  ylab(expression(paste("Biofilm thickness in ", mu, "m"))) +  plot.options.legend +  theme(axis.text.x = element_text(angle = 270), legend.title = element_blank())  ggplot(summary) +  geom_boxplot(aes(x = electrode, y = thickness), width = 0.2) +  scale_y_continuous(limits = c(0, 50), breaks = seq(0, 50, 10)) +  ylab(expression(paste("Biofilm thickness in ", mu, "m"))) +  xlab("") +  plot.options  # Statistical indicators are calculated as mean and SD of biofilm thickness, first for each z-image, then for each biofilm. The SD in the summary per biofilm is calculated as error propagation from the variances (= SD^2^) of the z-image summary.  # Plotting the thickness calculated for each z-image allows to detect outliers and to visually compare them to the actual image  summary_z 🡨 summary%>%  group_by(file, electrode)%>%  summarise(mean_1 = round(mean(thickness), 1), sd_1 = round(sd(thickness), 1))  summary_biofilm 🡨 summary_z%>%  group_by(electrode)%>%  summarise(mean_sum = round(mean(mean_1), 1), sd_sum = round(sqrt(sum(sd_1^2)), 1)) |
| --- | --- |
